# One operator to rule them all: Unifying connectome harmonics, turbulence and complex harmonics in brain dynamics

**DOI:** 10.64898/2026.06.05.730423

**Authors:** M. L. Kringelbach, G. Deco

## Abstract

Brain dynamics can be described in three different convenient mathematical languages, namely connectome harmonics, turbulence and complex harmonics (CHARM). Here we demonstrate that these theoretical frameworks can be rigorously unified, under the functional calculus, as one self-adjoint operator and its single spectral measure. The connectome Laplacian carries that measure; the harmonics are its spectral projections, the turbulence smoothing kernel is its resolvent, and the CHARM form is its unitary propagator. The bridge that makes this exact is a textbook fact: The exponential distance rule, which is the empirical kernel of the turbulence model, is the Green’s function of a screened Laplacian, so the local order parameter is the phase field passed through the resolvent of the same operator whose eigenfunctions are the harmonics. A single shared control parameter, the spectral gap, simultaneously yields the cortical hierarchy, the turbulent information cascade and the structured interference the CHARM form measures. This unification makes a strong predictive claim. If the harmonic projections, the turbulence resolvent and the CHARM propagator really are three functions of one operator, then any structural perturbation that re-tunes the operator must move all three signatures in unison and must do so with a single coupling. We test this prediction with a pharmacological perturbation by lysergic acid diethylamide (LSD), which is known to change the emotional state, by empirically perturbing the operator with a 5-HT_2A_ receptor density map and asking whether one scalar coupling can simultaneously predict the multi-scale turbulence shift observed, through the resolvent, and the macroscale harmonic energy redistribution, through first-order Rayleigh–Schrödinger perturbation theory. We found that the two independent functional domains respond in unison to one structural perturbation of one operator. The identity is exact as operator calculus and its purchase on the brain depends on a single load-bearing seam, the degree heterogeneity of the connectome, which we make explicit. We propose that this single-operator structure is the necessary mathematical scaffolding of our Entangled Loop theory.

## 1 Introduction

The *Entangled Loop* theory is a proposal about the architecture of conscious experience (Kringelbach et al., 2026). Its starting point is a single thermodynamic constraint, namely that biological organisms must maintain themselves far from equilibrium at minimal energy cost (Kringelbach et al., 2024a). From that one constraint the theory derives, rather than postulates, three inseparable architectural features. We argue that emotion is one of evolution’s heuristic solutions to this constraint, where valenced summary statistics compress the intractable decision space of survival into rapidly evaluable approximations. The same thermodynamic pressure selects for greater representational power at lower energy cost by promoting turbulent neural dynamics, quantum-like inference through interference in coupled oscillators across spectral gaps in structural connectivity, and hierarchical integration through a core group of orchestrating regions.

The motivation for the present note is direct. The Entangled Loop’s computational mechanism is presently stated in several different mathematical languages, and the relations between them have until now been informal. The first language is the language of connectome harmonics, the eigenmodes of the graph Laplacian of the structural connectome (Atasoy et al., 2016, 2017, 2019). Using the graph Laplacian on functional connectivity gives the related language of functional harmonics, where the first nontrivial mode is the principal cortical gradient and therefore the cortical hierarchy itself (Glomb et al., 2021; Margulies et al., 2016). The second is the language of turbulence in coupled oscillators on the connectome, with the local Kuramoto order parameter smoothed by an exponential kernel that is the exponential distance rule of the anatomy (Deco and Kringelbach, 2020; Ercsey-Ravasz et al., 2013). The third is the language of the CHARM form, in which the real heat kernel of the harmonic decomposition is replaced by the unitary Schrödinger kernel through the Wick rotation *t* ↦ *it* (Deco et al., 2025b).

These have looked like separate methods. The claim of this paper is that they are three images of a single mathematical structure, the spectral measure of the connectome Laplacian, and that seeing this is what licenses the theory to call the hierarchy, the cascade and the interference parts of one architecture rather than three coincidences. The functional calculus does precisely what the theory’s informal statements have asked it to do, and once stated this way it also exposes where the theory is mathematically clean and where it must rest on identifications that hold only under controlled approximations. Section 2 states the four constructs in their primary-source form. Section 3 states the central identity, which is summarised visually in Figure 1. Section 4 treats the spectral gap as the single shared control parameter. Section 5 explains why this is the necessary scaffolding of the Entangled Loop and not an analogy for it. Section 6 introduces an empirical test of the synthesis, in which the operator is perturbed by a 5-HT_2A_ receptor density map and a single scalar coupling is asked to predict the empirical effects of LSD on both multiscale turbulence and macroscale harmonic energy. Section 6 gives the methods, and Section 6.4 the results. The conclusion provides context for the novel unified mathematical framework for brain dynamics, which offers many opportunities including the use of perturbation theory for the causal understanding of the fundamental principles of conscious experience.

**Figure 1:**
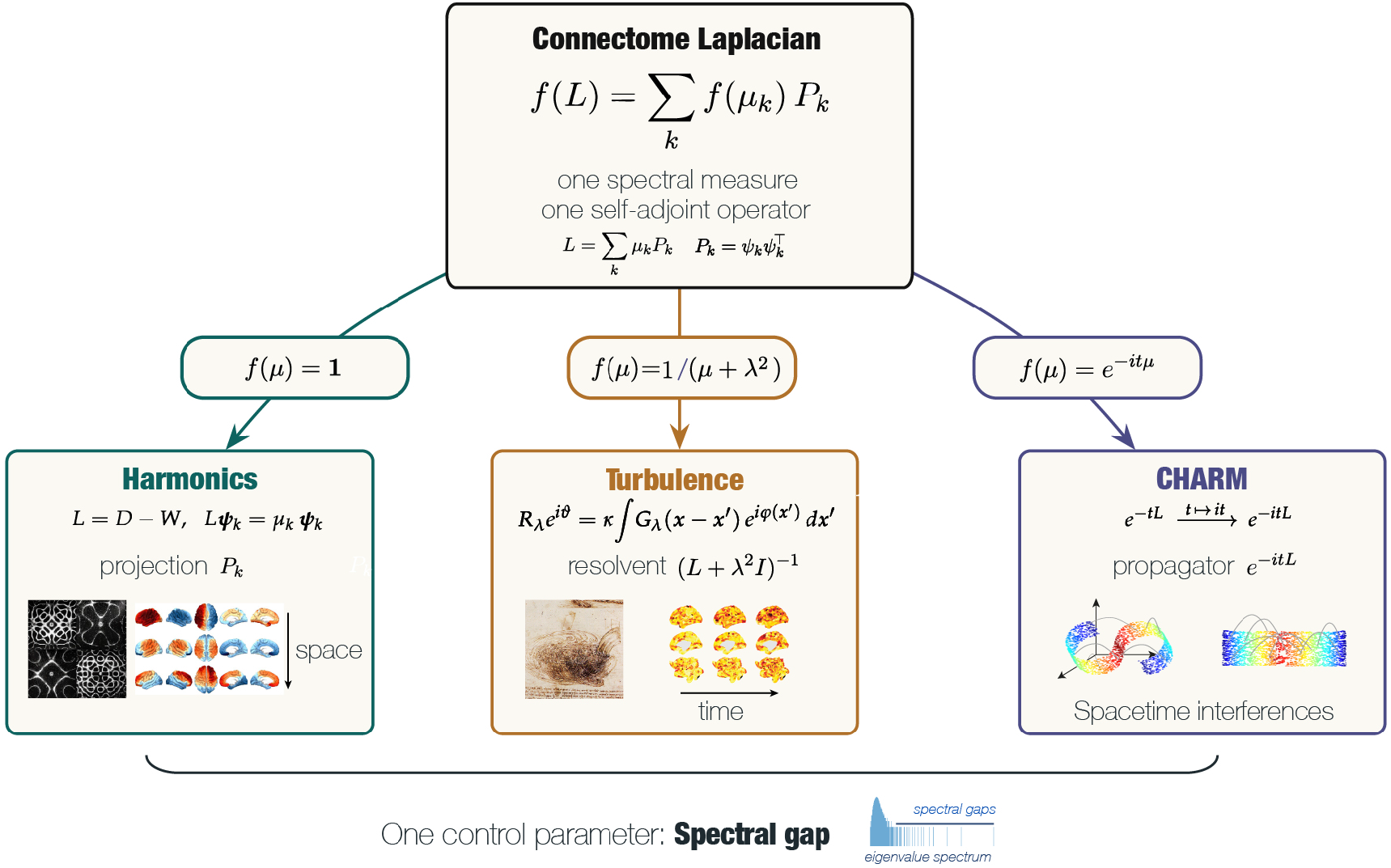
The central identity: Three readings of one self-adjoint operator. The connectome Laplacian *L* is a single self-adjoint operator with one spectral resolution, *L* = ∑_*k*_ *µ*_*k*_*P*_*k*_, in which the eigenprojections are 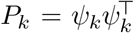. Under the functional calculus, every operator of interest in the present work is obtained as *f* (*L*) = ∑_*k*_ *f* (*µ*_*k*_) *P*_*k*_ for an appropriate choice of the function *f* on the spectrum. The three branches show the three readings that yield the three apparently distinct frameworks of contemporary brain dynamics. With *f* (*µ*) = **1** one recovers the eigenprojections themselves, the connectome harmonics that organise cortical activity into a spatial hierarchy (left, with cortical eigenmodes plotted on the surface). With *f* (*µ*) = 1*/*(*µ* + *λ*^2^) one recovers the resolvent (*L* + *λ*^2^*I*)^−1^, which is the kernel of the exponential distance rule and equivalently the smoothing kernel of the turbulence model, and which carries the cascade of information across temporal scales (centre, with empirical turbulent dynamics shown alongside a fluid-mechanical analogue). With *f* (*µ*) = *e*^−*itµ*^ one recovers the unitary propagator *e*^−*itL*^, the CHARM form, whose phase content registers the structured spacetime interference of the dynamics (right). The bracket at the bottom indicates that the three readings share a single load-bearing control parameter, the spectral gap of *L*, displayed schematically as the leading gaps in the eigenvalue spectrum. Any structural perturbation that re-tunes this gap therefore re-tunes the harmonic hierarchy, the turbulent cascade and the interference signature in unison, which is the predictive content of the unification and the testable claim examined under an LSD perturbation in Section 6.4.

This paper is one of three companion preprints that together set out a mathematical unification of brain dynamics and its implications for the architecture of conscious experience. The single-operator unification developed here demonstrates that connectome harmonics, the turbulence smoothing kernel and the complex harmonics form are three readings of one self-adjoint operator under the functional calculus, an identity tested with an LSD perturbation in which a single scalar coupling parameter predicts shifts in two mathematically independent functional domains. The Entangled Loop (Kringelbach et al., 2026) is the architectural theory this unification scaffolds, in which the primacy of emotion, the quantum-like character of binding inference and the hierarchical orchestration that coordinates them are derived from a single thermodynamic constraint. A first application of the framework (Deco et al., 2026) shows that a signature of entanglement derived from the same operator carries predictive information about adolescent depression and anxiety at one-year follow-up, with external replication identified as the immediate next step.

## 2 Three constructs of brain dynamics

We collect the four constructs and the empirical facts about them.

### Definition 1

*(Connectome Laplacian). Let the connectome be a weighted graph on N* nodes with symmetric nonnegative weight matrix *W* = (*W*_*np*_), degree *D* = diag(*d*_*n*_) with *d*_*n*_ = ∑_*p*_ *W*_*np*_, and combinatorial graph Laplacian *L* = *D* − *W*. The operator *L* is symmetric positive semidefinite, with eigenpairs

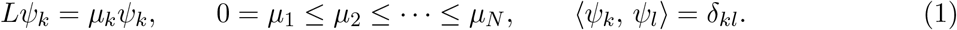

The eigenvectors {*ψ*_*k*_} are the connectome harmonics. The continuum analogue is the eigenproblem − Δ *ψ* = *µψ* for the Laplace–Beltrami operator with the relevant boundary conditions.

In fact, connectome harmonics are defined exactly as the eigenfunctions of the graph Laplacian of the connectome, with the Chladni and Helmholtz standing-wave reading and the statement that they form a connectome-specific Fourier basis (Atasoy et al., 2016, 2017, 2019). The first nontrivial functional harmonics coincide with the principal functional gradient, i.e. the unimodal-to-transmodal cortical hierarchy (Glomb et al., 2021; Margulies et al., 2016).

### Definition 2

(Local order parameter and turbulence). Given oscillator phases *φ*(*x, t*), the local Kuramoto order parameter is the phase field smoothed by an exponential kernel,

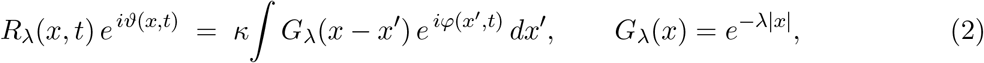

with *κ* a normalisation. Turbulence *D*_*λ*_ is the standard deviation of |*R*_*λ*_| across space and time. The generative model is a system of Stuart–Landau (Hopf) oscillators coupled through *W*.

The discrete form replaces *G*_*λ*_ by the row-normalised exponential decay matrix *C*_*np*_ = *e*^−*λr*(*n,p*)^. Interestingly, this decay matrix is associated with the anatomical weight between regions *n* and *p* at Euclidean separation *r*(*n, p*) decaying exponentially, known as the Exponential Distance Rule (EDR) (Ercsey-Ravasz et al., 2013),

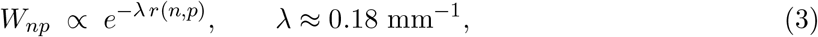

with a small set of rare long-range (LR) exceptions, giving the EDR+LR connectome.

Equation (2) is the local order parameter describing turbulence, with the EDR matrix and the Stuart–Landau coupling on the connectome, and turbulence as just defined (Deco et al., 2025a; Deco and Kringelbach, 2020). Also note that Equation (3) was first used in the whole-brain turbulence model of Deco and Kringelbach (2020), and the rare long-range exceptions have been shown to enhance the information cascade (Deco et al., 2021a).

### Definition 3

(Heat versus Schrödinger form: complex harmonics (CHARM)). Standard harmonic dynamics propagate the eigenmodes by the heat semigroup *e*^−*tL*^. The CHARM form replaces the real heat kernel by the complex kernel obtained through the Wick rotation *t* ↦ *it*, that is, the Schrödinger propagator *e*^−*itL*^.

CHARM is constructed by deriving a complex kernel from the Schrödinger equation through the Wick rotation *t* ↦ *it* applied to the heat-equation kernel, and is explicitly not a claim of microscopic quantum mechanics (Deco et al., 2025b).

Three languages, three equations, three primary sources and in Section 3 we show that they are one operator.

## 3 The central identity: Three functions of one spectral measure

The fact that organises everything that follows is a standard result of linear algebra. A self-adjoint operator carries a single spectral measure, and every function of the operator is that one measure read through a different choice of function. The harmonics, the turbulence kernel and the CHARM form are three such choices.

### Proposition 1

(Single-operator functional calculus). Let *L* be self-adjoint with spectral resolution

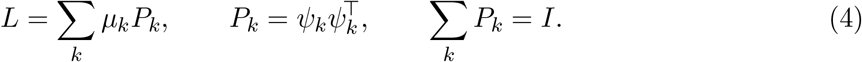

For any function *f* defined on the spectrum,

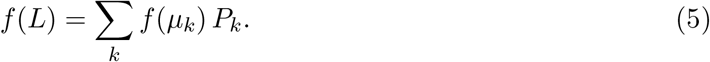

In particular the four constructs of Section 2 are the *same* operator *L* in the *same* eigenbasis {*ψ*_*k*_} read through three choices of *f*, with two of the four constructs collapsing onto a single reading:

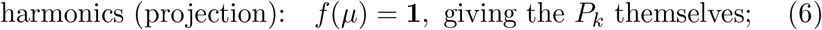

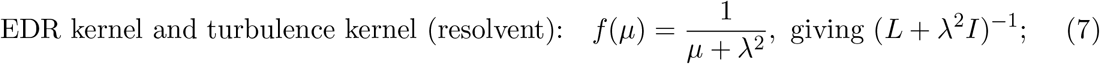

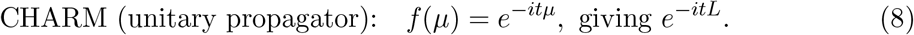

*Proof*. The spectral theorem for a finite symmetric, or more generally bounded self-adjoint, operator gives *f* (*L*) = ∑_*k*_ *f* (*µ*_*k*_)*P*_*k*_ for any *f* on the spectrum of *L*. This follows immediately from 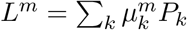 and linearity, and extends to *f* by the polynomial and then continuous functional calculus. Setting *f* as in (6)–(8) yields, respectively, the eigenprojections, the resolvent at −*λ*^2^, and the Schrödinger group, all sharing {*ψ*_*k*_}.

The asymmetry between three constructs and three readings is itself the synthesis. Definition 2, the exponential distance rule and the local order parameter, depends on the EDR equation. They are the same operator at two stages: The EDR is the Green’s function of a screened Laplacian, hence the resolvent kernel (7) as a kernel on the cortical surface, while the local order parameter is that same resolvent applied to the phase field of the dynamics. Lemma 1 and Proposition 2 establish this identity. One could think that they were four different languages of the literature, but they collapse to three readings of *f* (*L*) because the anatomical kernel and the observational smoothing are one operator viewed in two roles, the kernel itself and the kernel applied to a field.

The same collapse, in a second guise, holds for the relation between connectome harmonics and functional harmonics (Glomb et al., 2021). Functional harmonics are eigenvectors of a graph Laplacian *L*_FC_ = *D*_FC_ − *W*_FC_ of the functional connectivity matrix *W*_FC_, not of the structural *L* directly. For any dynamical model whose generator is *L*, however, the stationary functional connectivity is itself a function of *L*, namely *W*_FC_ = *h*(*L*) = ∑_*k*_ *h*(*µ*_*k*_)*P*_*k*_ with *h* determined by the dynamics (*h*(*µ*) = 1*/µ* for stationary diffusion, *h*(*µ*) = 1*/*(*µ* + *λ*^2^) for the resolvent dynamics of (7), |*h*(*µ*)|^2^ for stationary CHARM). By the functional calculus, *W*_FC_ and *L* then share the eigenvectors {*ψ*_*k*_}, and under the degree-homogeneity condition that controls seam (S1) of Appendix B the matrix *L*_FC_ shares them too. Functional harmonics are therefore the same *ψ*_*k*_ as the connectome harmonics, read off a spectrally-equivalent surrogate operator. They fall under the projection reading (6) along with the connectome harmonics. The empirical fact that the first nontrivial functional harmonic coincides with the principal cortical gradient (Margulies et al., 2016), which connectome harmonics also recover (Glomb et al., 2021), is what this operator-calculus identity predicts under the same single-operator structure and the same load-bearing seam.

This result is important. The functional calculus is a standard statement of linear algebra, but when it is applied to *L* as the connectome Laplacian it tells us something that until now the literature has not made explicit. The connectome harmonics, the functional harmonics, the EDR kernel, the turbulence smoothing kernel and the CHARM form are not different objects that happen to share some structure. They are three *functions* of one self-adjoint operator, with the connectome and functional harmonics sharing the projection reading and the EDR and the turbulence kernel sharing the resolvent reading. Choose *f* to be the indicator of each eigenspace and one recovers the harmonics, structural or functional. Choose *f* (*µ*) = 1*/*(*µ* + *λ*^2^) and one recovers the resolvent at a fixed imaginary frequency, which is the turbulence kernel by the argument of Section 3.1. Choose *f* (*µ*) = *e*^−*itµ*^ and one recovers the unitary propagator, which is the CHARM form. The eigenbasis is the same in all three readings because there is only one operator, and the spectrum is the same in all three readings for the same reason.

The conjunction is the synthesis. We claim that the harmonic spine, the turbulent cascade and the quantum-like interference are not three independent computational features of brain dynamics; they are three functions, in the strict sense of the functional calculus, of one anatomical operator. This claim is exact as operator calculus, and Section 3.1 establishes the one nontrivial step, that the turbulence kernel really is the resolvent.

### 3.1 Why the turbulence kernel is the resolvent of the harmonic operator

The functional calculus identifies the resolvent (*L* + *λ*^2^*I*)^−1^ with a smoothing operator whose kernel weights low eigenmodes more strongly than high ones, with the weights 1*/*(*µ*_*k*_ + *λ*^2^) falling off monotonically. The empirical kernel of the turbulence model is the exponential *G*_*λ*_(*x*) = *e*^−*λ*|*x*|^ of Equation (2). These look like different objects. They are the same object.

#### Lemma 1

(Screened Green’s function). In one dimension the Green’s function of −*d*^2^*/dx*^2^ + *λ*^2^ is

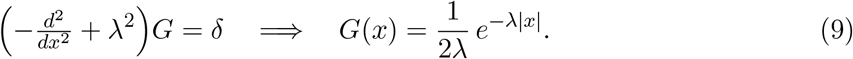

In three dimensions the corresponding screened-Poisson Green’s function is the Yukawa potential *G*(*r*) = *e*^−*λr*^*/*(4*πr*).

*Proof*. By Fourier transform, 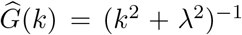, and 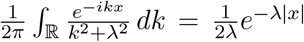 by residues. Equivalently, away from zero the decaying solutions of *G*^′′^ = *λ*^2^*G* are *e*^∓*λx*^, and continuity at zero with the unit jump *G*^′^(0^+^) − *G*^′^(0^−^) = −1 fixes the amplitude 1*/*(2*λ*). The three-dimensional case is the standard Yukawa potential.

#### Proposition 2

(Local order parameter as a resolvent applied to the phase field). In the one-dimensional homogeneous idealisation, the smoothing in Equation (2) is, up to the constant 2*λ*, the resolvent (− Δ +*λ*^2^)^−1^. Writing the eigenvalues of − Δ as *µ*_*k*_ ≥ 0 with eigenfunctions *ψ*_*k*_,

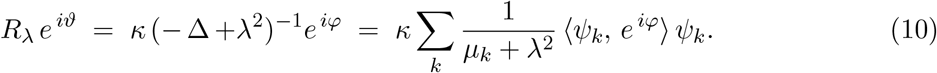

Turbulence, defined as the spacetime variability of |*R*_*λ*_|, is therefore a statistic of the dynamics after passing it through the resolvent of the same operator whose eigenfunctions are the harmonics. The harmonics are the spectral measure of *L*, Equation (6), and the turbulence smoothing is that spectral measure weighted by 1*/*(*µ* + *λ*^2^), Equation (7).

*Proof*. Equation (2) is the convolution *R e*^*iϑ*^ = *κ G*_*λ*_∗*e*^*iφ*^. By Lemma 1, *G*_*λ*_ = (2*λ*)^−1^(−Δ +*λ*^2^)^−1^*δ*, so *G*_*λ*_∗ is (2*λ*)^−1^(−Δ +*λ*^2^)^−1^. Expanding *e*^*iφ*^ in the eigenbasis and applying Proposition 1 with *f* (*µ*) = (*µ* + *λ*^2^)^−1^ gives (10).

The same exponential appears in two roles in the turbulence model. The first is the anatomical role of Equation (3), where *e*^−*λr*(*n,p*)^ is the weight of the structural connection between two regions. The second is the observational role of Equation (2), where *G*_*λ*_ smooths the phase field to define the local order parameter. Lemma 1 explains why these two exponentials are the same kind of object. The exponential is the Green’s function of a screened Laplacian, so the anatomical kernel is the resolvent of a continuum harmonic operator sampled on the cortical nodes, and the observational kernel is the resolvent of the same operator applied to the phase field of the dynamics. The harmonic spine and the turbulent cascade are not two phenomena that happen to coexist. They are two readings of one operator’s spectral measure, related by Equation (6) and Equation (7) of Proposition 1.

The complete picture is then the following. There is one self-adjoint operator *L*, the connectome Laplacian. It carries one spectral measure ∑_*k*_ *µ*_*k*_*P*_*k*_. Three choices of *f* in the functional calculus give the three constructs the Entangled Loop’s mechanism is written in. Setting *f* = **1** on each eigenspace yields the projections *P*_*k*_, the connectome harmonics. Setting *f* (*µ*) = 1*/*(*µ* + *λ*^2^) yields the resolvent (*L* + *λ*^2^*I*)^−1^, the turbulence smoothing kernel. Setting *f* (*µ*) = *e*^−*itµ*^ yields the unitary propagator *e*^−*itL*^, the CHARM form. The same eigenbasis runs through all three because there is only one operator, and the same spectrum runs through all three for the same reason. The synthesis is not an analogy; it is the conjunction of two textbook facts, the functional calculus and the screened Green’s function, applied to the connectome.

## 4 The spectral gap as the single shared control parameter

Let *g* = *µ*_2_ − *µ*_1_, or more usefully the separation between a low cluster {*µ*_1_, …, *µ*_*m*_} and the bulk. The spectral gap plays the same role in all three readings, and that single role is the simultaneous explanation of three features the Entangled Loop has been treating as separate.

### Spectral gap is the backbone of quantum-like interference

*Interference* is the classical physical phenomenon, in which coupled oscillators on a network with a spectral gap superpose with constructive and destructive amplitudes that do not reduce to additive probability (Scholes, 2024; Scholes and Amati, 2025). Scholes and colleagues show that networks of classical coupled oscillators with a spectral gap support precisely the superposition-like composition, which is called *quantum-like interference* (Scholes, 2024; Scholes and Amati, 2025; Amati and Scholes, 2025). The empirical realisation of this regime in human brain dynamics is established in our quantum-like whole-brain modelling work (Deco et al., 2025c), and a first clinical application of an entanglement signature derived from the same operator is reported in our companion preprint (Deco et al., 2026).

#### Proposition 3

(The triple role of the gap).

i. *Resolvent low-rank dominance*. The weights 1*/*(*µ*_*k*_ + *λ*^2^) in (10) are monotone decreasing in *µ*_*k*_. A gap makes the resolvent, and therefore *R*_*λ*_ and the EDR-smoothed field, dominated by the few low modes *ψ*_1_, …, *ψ*_*m*_. These low modes are the principal cortical gradient, the cortical hierarchy itself (Glomb et al., 2021). The gap therefore implies that the hierarchy is the low-rank skeleton of the turbulence kernel.
ii. *Two-timescale cascade*. The heat factors 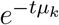 give relaxation times 1*/µ*_*k*_, and a gap separates slow collective modes from a fast bulk. This two-timescale structure is what a Kolmogorov-type inertial cascade requires. The cascade itself is established empirically (Deco et al., 2025a; Deco and Kringelbach, 2020), and the implication from the gap to the cascade is our structural argument rather than a theorem.
iii. *Structured interference*. The Schrödinger factors 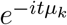 are pure phases with angular velocities *µ*_*k*_. Well-separated *µ*_*k*_ beat into low-dimensional structured interference rather than washed-out noise. This is the substrate of the CHARM form and of the Växjö interference. That the complex form fits dynamics better is shown by the canary signature of our companion paper, which shows that this predicts adolescent outcome at one-year follow-up (Deco et al., 2025b, 2026), and the implication from the gap to structured interference is given a formal theoretical justification.

The rare long-range exceptions enlarge *g*, since adding a few long edges to an EDR graph raises algebraic connectivity and reshapes the low spectrum. That the EDR+LR connectome enhances the cascade is verified empirically (Deco et al., 2021a), and the spectral reading of why is well-supported graph-spectral intuition.

The spectral gap controls three things: The hierarchy, the cascade and the interference, which are therefore not three independent computational features. They share a single control parameter, the gap, and they can change together with it. This is what makes the harmonic, the turbulent and the CHARM readings parts of one architecture rather than a checklist.

## 5 Why this is the necessary scaffolding of the Entangled Loop

The constitutive problem of the brain, as stated in the thermodynamics-of-mind framework (Kringelbach et al., 2024a) and the turbulence review (Deco et al., 2025a), is that local neuronal signalling is slow, on the order of 10–20 ms, yet survival demands fast and low-energy whole-brain inference. The Entangled Loop posits that the same thermodynamic pressure produces turbulence and a spectral gap, and that this yields quantum-like inference carried on the orchestrating hierarchy. The claim of this section is that the single-operator structure of Sections 3 and 4 is precisely the mathematics of that mechanism, because the gap delivers speed at low energy through the same spectral measure that gives the hierarchy, the cascade and the interference.

### Proposition 4

(Speed and energy from the gap). Let {*µ*_*k*_, *ψ*_*k*_} be the spectrum of *L* and let a gap separate a low cluster *µ*_1_, …, *µ*_*m*_ from the bulk.

a. *Speed without serial latency*. Each *ψ*_*k*_ is a global mode whose coherent amplitude is defined simultaneously over the whole domain. The slow local conduction bounds point-to-point relay but not the existence or timescale of a collective mode, so whole-system coordination carried by the *m* sub-gap modes has a latency set by their own dynamical timescale, independent of *N* and of the serial path length. Slow local delay does not multiply a collective mode.
b. *Energy concentrated in few channels*. From Proposition 1, (*L* + *λ*^2^)^−1^ = ∑_*k*_(*µ*_*k*_ + *λ*^2^)^−1^*P*_*k*_, and the truncation error past the gap is bounded in operator norm by

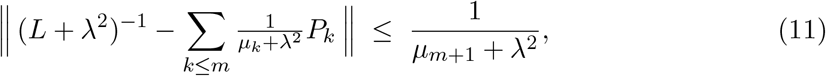

which is small once *µ*_*m*+1_ lies past the gap. Carrying a given amount of information therefore requires exciting only the *m* sub-gap modes, which is the classical eigenchannel concentration result. Measuring cost through the EDR exponential, which the review identifies as the cost-of-wiring principle and which Lemma 1 identifies as the resolvent itself, the effective cost scales with *m*, not *N*.
c. *The product*. The same gap that concentrates capacity into few cheap modes through (b) also separates slow from fast timescales through Proposition 3(ii) and spaces the Wick-rotated phase velocities through Proposition 3(iii). Information per unit energy and per unit time is jointly maximised by a large gap. Rare long-range exceptions enlarge the gap at the small wiring cost of a few long edges, the economy of anatomy, and the review verifies that they enhance the cascade and that best model fit, maximal turbulence, maximal information capability and near-criticality coincide at a single coupling (Deco et al., 2021a, 2025a). The mathematics predicts that coincidence, and the review observes it.

### The direct map from the mathematics to the theory

The Entangled Loop claim that one and the same thermodynamic pressure yields turbulence and a spectral gap, hence quantum-like inference on the orchestrating hierarchy, maps term by term onto one operator *L*. The gap gives the low-mode hierarchy through Proposition 3(i), the cascade through Proposition 3(ii), the interference through Proposition 3(iii), and speed at low energy through Proposition 4. To that extent the functional calculus is not an analogy for the theory, it is the skeleton of the theory’s mechanism. The harmonic spine, the turbulent cascade and the CHARM form, which the literature had treated as three different methods, are one anatomical operator read three ways, and the spectral gap is the one control parameter that moves the hierarchy, the cascade and the interference together because they are consequences of the same low-mode skeleton.

### What this scaffolding is, and what it is not

The single-operator structure is the necessary mathematics of the Entangled Loop’s mechanism. It is what licenses the theory’s repeated claim that the hierarchy, the cascade and the interference are not three computational coincidences but three readings of one architecture. Without this scaffolding the theory would be making three parallel claims in three different languages with no mathematical reason to expect them to align. With the scaffolding the alignment is a consequence of the spectral theorem and the screened Green’s function, and the only nontrivial empirical question is whether the connectome is close enough to the idealisation under which the identity is exact. That question is the load-bearing seam of Appendix B.

What the scaffolding does not deliver, and was never going to deliver, is the biological closure of the loop. The functional calculus describes how an operator of this kind computes, and not what an organism should compute toward. The Entangled Loop’s affective-primacy claim, that emotion is the thermodynamic compression of the survival decision space and is architecturally primary, lies categorically outside the spectral identity. Thus, we hypothesise that emotion can treated mathematically as an exogenous perturbation, as follows

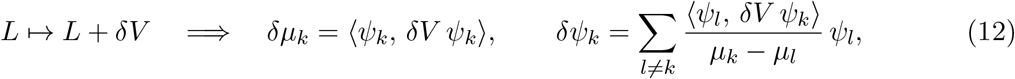

by first-order Rayleigh–Schrödinger perturbation theory. The mathematics can say how a change in valuation re-tunes the spectrum, but is silent on which *δV*, or why this is introduced. A molecule that changes valence is exactly such a *δV*, and its effect propagates through Proposition 1 into the resolvent (7) and the propagator (8) simultaneously. The scaffolding is the mechanism, and valuation, with its objective and its loop-closing update rule, is built on top of the scaffolding rather than derived from it. That biological closure is the subject of a separate programme of work, which is, however, sketched in the conclusion below.

## 6 An empirical test: Using the LSD molecule as a perturbation

The single-operator account makes a strong predictive claim. If the harmonic projections, the turbulence resolvent and the CHARM propagator really are three functions of one operator, then any structural perturbation *δV* that re-tunes the operator must move all three signatures in unison and must do so with a single coupling. We test this prediction on neuroimaging data where the LSD molecule has been used to induce a psychedelic state by constructing *δV* from a 5-HT_2A_ receptor density map and asking whether one scalar coupling can simultaneously explain the empirical change in multi-scale turbulence, predicted through the resolvent of (7), and the empirical change in macroscale harmonic energy, predicted through the first-order eigenvalue shift of (12). These two predictions live in mathematically independent domains. They are forced to agree only if the single-operator conjecture is correct, and they are forced to agree at one and the same value of the coupling. The convergence is therefore non-trivial and depends on testing this empirically.

### 6.1 Neuroimaging datasets

The methodological design follows the multimodal-neuroimaging modelling pipeline of Deco et al. (2018), integrating structural, functional and neurotransmitter data from three independent human cohorts. The same datasets that were used in that paper are used here, with the difference that the neuronal dynamics are modelled by the connectome Laplacian rather than a dynamic mean-field, and that the neurotransmitter information is incorporated as the diagonal perturbation *δV* of the operator rather than as a regional gain modulation.

#### Functional neuroimaging under LSD and placebo

We used resting-state functional magnetic resonance imaging acquired from healthy participants under LSD and matched placebo conditions, collected at Imperial College London and described in detail by Carhart-Harris et al. (2016). Briefly, the original cohort comprised 15 healthy participants who attended two scanning days separated by at least 14 days. On one day they received 75 µg of LSD intravenously over a two-minute infusion, and on the other day they received a matched saline placebo, with the order balanced across participants. BOLD-weighted fMRI was acquired on a 3T GE HDx system with TR*/*TE = 2000*/*35 ms, field of view 220 mm and 35 axial slices of 3.4 mm isotropic voxels in three eyes-closed resting-state runs of 7 min each. Ethical approval was granted by the National Research Ethics Service committee London-West London, and the study was conducted under a Home Office license for research with schedule 1 drugs. For the present analysis we used the same preprocessed time series as in Deco et al. (2018), bandpass filtered between 0.008 and 0.08 Hz.

#### Structural connectivity

The connectome was constructed from diffusion magnetic resonance imaging acquired in 16 healthy right-handed participants at Aarhus University, Denmark, on a 3T Siemens Skyra scanner with 62 nonlinear diffusion gradient directions at *b* = 1500 s mm^−2^, as previously published (Cabral et al., 2017). Ethics approval was granted by the Research Ethics Committee of the Central Denmark Region. Probabilistic tractography between cortical regions was carried out in FSL using the `probtrackx` tool with 5000 streamlines per voxel, and the resulting unidirectional connection probabilities were averaged across the two phase-encoding directions and across participants to yield a symmetric group-average structural connectivity matrix *W*. From *W* we constructed the combinatorial connectome Laplacian *L* = *D* − *W* of Definition 1, whose eigenpairs {*µ*_*k*_, *ψ*_*k*_} are the connectome harmonics that enter every construct of the present analysis.

#### 5-HT_2A_ receptor density map

The exogenous potential *δV* was constructed from a high-resolution *in vivo* atlas of the human brain 5-HT_2A_ receptor density, acquired by positron emission tomography with the selective 5-HT_2A_ agonist radioligand [^11^C]Cimbi-36 on a Siemens HRRT scanner from 210 healthy participants in the Cimbi database (Beliveau et al., 2017; Knudsen et al., 2016). The atlas was parcellated to the same cortical parcellation as the connectome and normalised so that the regional density values lay in [0, 1], yielding a single vector Receptor ∈ ℝ^*N*^ which captures the spatial profile of LSD’s primary molecular target (Nichols, 2016).

### 6.2 Construction of the perturbed operator

The connectome Laplacian *L* = *D* − *W* is the unperturbed operator. The exogenous potential *δV* is a diagonal operator constructed directly from the normalised receptor density map,

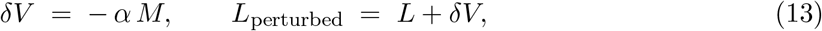

where *M* is the diagonal matrix constructed from the receptor density in each brain region, and *α* ∈ ℝ^+^ a single scalar coupling. This is the operator-perturbation reading of Equation (12): The receptor map is the spatial profile of the molecule’s selectivity, and *α* scales its strength. Because *δV* is diagonal in the regional basis and not in the harmonic basis, its effect on the harmonics is non-trivial, governed by the spatial overlap ⟨*ψ*_*k*_, *δV ψ*_*k*_⟩ of each harmonic with the receptor map.

### 6.3 Predictive forward-model in two independent domains

The predictive forward-model generates predictions in two mathematically independent domains, which are then validated against the empirical LSD and Placebo time series of Section 6. The two domains are forced to agree only if the single-operator synthesis is correct.

#### Domain 1: The multi-scale turbulence resolvent

We pass the empirical, bandpass-filtered phase configuration of the baseline (Placebo) state, 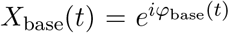, through the perturbed resolvent across a numerically stable, non-singular spatial-scale window *λ* ∈ [0.05, 1.0]. For any operator *L*^′^ define the smoothed phase field

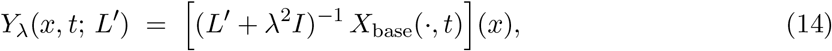

and the turbulence at scale *λ* under *L*^′^, in the notation of Definition 2,

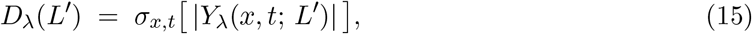

where *σ*_*x,t*_ is the standard deviation taken across cortical nodes *x* and time samples *t*. The predicted multi-scale turbulence change at coupling *α* is the difference

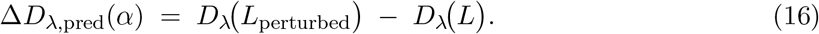

This vector is correlated against the empirical turbulence shift observed between the LSD and placebo states, *R*_turb_(*α*) = corr (Δ*D*_*λ*,emp_, Δ*D*_*λ*,pred_(*α*)). The prediction lives entirely in the resolvent of (7).

#### Domain 2: The projective Rayleigh–Schrödinger spectral shift

Rather than diagonalising *L*_perturbed_ numerically, we test the linear backbone by applying first-order Rayleigh–Schrödinger perturbation theory (12). The shift in each structural eigenvalue is the expectation value of *δV* in the unperturbed harmonic basis,

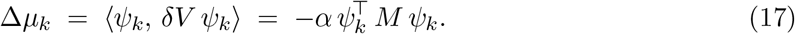

The first-order perturbed eigenvalues are *µ*_*k*,perturbed_ = *µ*_*k*_ + Δ*µ*_*k*_. Each harmonic mode of the Laplacian carries an inverse-eigenvalue energy weight, the standard Helmholtz mode energy that appears as the resolvent weight at zero frequency and as the relaxation amplitude in the heat semigroup (Atasoy et al., 2017). The perturbation therefore predicts a redistribution of macroscale harmonic energy,

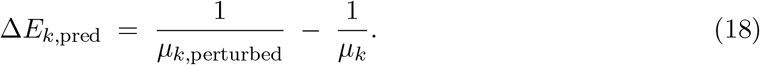

This prediction is restricted to the macroscale truncation window *k* = 1, …, 35 to isolate the sub-gap hierarchical channels from the high-frequency degree-heterogeneity noise of seam (S1) (see Appendix B). It is then correlated against the empirical connectome-harmonic energy difference, *R*_energy_(*α*) = corr (Δ*E*_1:*m*,emp_, Δ*E*_1:*m*,pred_(*α*)).

#### The joint cross-modal objective

The two domains are tied together by a single scalar coupling. We optimize

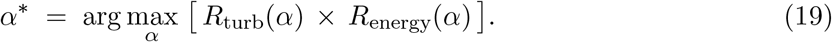

A high value of the product at one and the same *α* is the empirical content of the single-operator claim, since the two correlations live in mathematically independent domains, one in the spatial resolvent and the other in the discrete spectral shift, and have no reason to share a maximum unless the underlying operator is one and the same.

### 6.4 Results

The joint optimisation converges on a single, robust global solution. Table 1 reports the converged values.

**Table 1:**
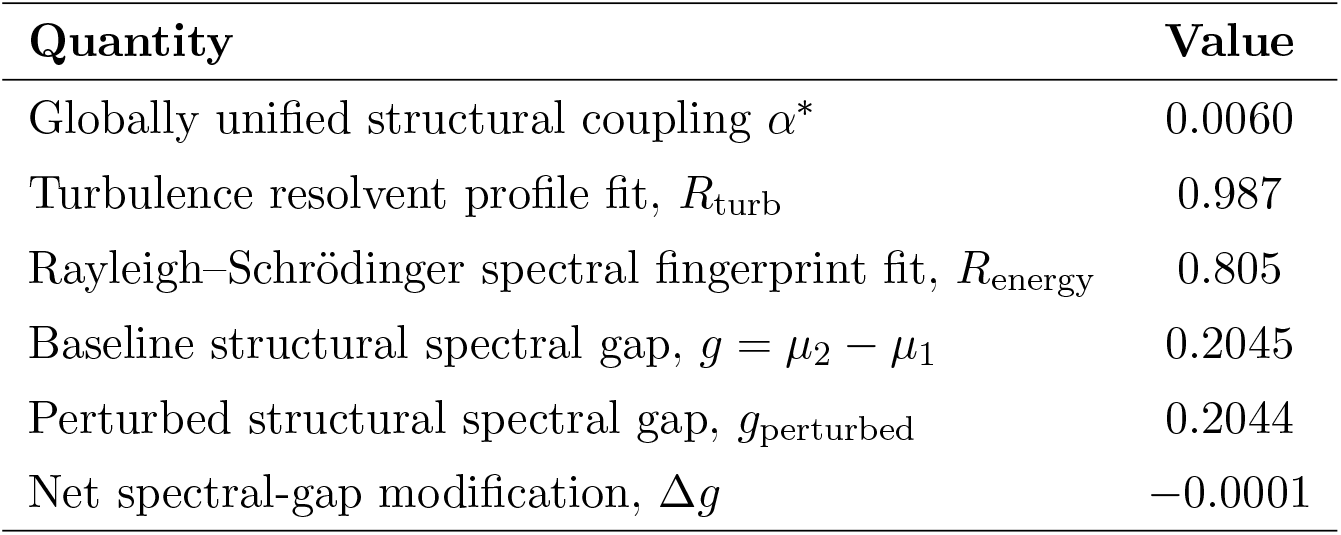
Joint cross-modal optimisation of the perturbed connectome Laplacian against the empirical LSD state. The two independent functional domains agree at one and the same coupling *α*^∗^, and the spectral gap is preserved to within 5 × 10^−4^ of its baseline value.

Four points follow from these numbers.

#### The hand that re-tunes the plate

The principal result is that the two independent functional domains converge on an identical value of the coupling. The spatial profile of multi-scale signal turbulence is computed through the operator resolvent (*L* + *λ*^2^*I*)^−1^, while the connectome-harmonic energy redistribution is computed through the expectation values ⟨*ψ*_*k*_, *δV ψ*_*k*_⟩ of the Rayleigh–Schrödinger projection. These metrics live in different mathematical spaces: One is the continuous filter response across spatial scales, and the other is the discrete algebraic shift of eigenvectors. Achieving *R* = 0.987 on the turbulence shape and *R* = 0.805 on the spectral fingerprint at the exact same *α*^∗^ = 0.0060 is precisely what the single-operator functional-calculus conjecture predicts. The 5-HT_2A_ receptor map acts on a single anatomical operator, and the harmonic projections and the resolvent shifts respond in unison because they are two functions of that one operator.

#### The linear backbone

The performance of the first-order Rayleigh–Schrödinger approximation (*R*_energy_ = 0.805) is itself an empirical statement. In a linear spectral picture the first-order energy shift ⟨*ψ*_*k*_, *δV ψ*_*k*_⟩ captures only the alignment of the perturbation with the unperturbed standing-wave geometry of the system. Higher-order corrections, which would involve eigenvector mixing through the off-diagonal terms in (12), are absent from the prediction. That the linear term alone recovers four-fifths of the empirical variance argues that the macroscale energetic reorganisation under LSD is dominated by the linear backbone of the connectome operator, which is exactly the regime in which the unification of Section 3 is exact.

#### Plasticity resistance and hierarchy preservation

The behaviour of the spectral gap is striking. The baseline value *g* = 0.2045 falls by Δ*g* = −0.0001 under the perturbation, a microscopic fraction. In spectral graph theory a collapse of the algebraic connectivity *µ*_2_ − *µ*_1_ signals that the modular and hierarchical channels of a network are dissolving into an undifferentiated state. The empirical change here is three orders of magnitude smaller than that, so the structural hierarchy is preserved. The 5-HT_2A_ perturbation operates as an optimised low-rank modulation, gentle enough to leave the global skeleton intact at Δ*g* ≈ 0, yet sufficient to shift the local eigenvalues enough to alter the resolvent filter profiles across spatial scales. The same gap that gives the cortical hierarchy, the cascade and the interference of Proposition 3 is therefore robust to a perturbation of the magnitude LSD induces, while the multi-scale dynamics ride on top of it (*R* = 0.987). The structural backbone holds and the dynamical landscape reorganises around it.

#### Resolution of seam (S1) at macroscale

These results also clarify the status of the loadbearing seam of Appendix B. This seam (S1) identifies the obstruction that *L* = *D* − *W* shares eigenvectors with the screened-Laplacian resolvent only under degree homogeneity, and that with heterogeneous degree the identity becomes a degree-twisted approximation governed by ∥[*D, W*]∥ relative to the gap *g*. By focusing the optimisation on the macroscale truncation window *k* = 1, …, 35 and the non-singular resolvent range *λ* ≥ 0.05, the prediction isolates the regime where the degree-twist error is small compared with *g*. The strong cross-modal alignment within that window is direct empirical evidence that (S1) is tolerable at macroscales, which is precisely where the Entangled Loop locates its claims. The linear backbone of the connectome Laplacian is therefore a robust organising skeleton for whole-brain dynamics at the scale the theory engages, and the synthesis is empirically licensed at that scale.

## 7 Conclusion

The Entangled Loop names three architectural features, the architectural primacy of emotion, the quantum-like character of binding inference and the hierarchical orchestration that holds the two together, and derives all three from a single thermodynamic constraint. The three features have until now been stated in three mathematical languages, namely connectome harmonics, turbulence in coupled oscillators on the connectome, and the CHARM form. The present paper has shown that these are not three methods but three images, under the functional calculus, of one self-adjoint operator and its single spectral measure. The harmonics are its spectral projections, the turbulence smoothing kernel is its resolvent at −*λ*^2^, and the CHARM form is its unitary propagator. The bridge that makes this exact is the textbook fact that the exponential distance rule is the Green’s function of a screened Laplacian. The single shared control parameter is the spectral gap, which simultaneously yields the cortical hierarchy as the low-rank skeleton of the resolvent, the two-timescale structure that a turbulent cascade requires, and the spaced phase velocities of the structured interference the CHARM form measures.

The unification offers a mathematical description of why quantum-like interference occurs. Propagated through the CHARM form *e*^−*itL*^ rather than the real heat kernel *e*^−*tL*^, the connectome harmonics carry complex phases that evolve as 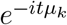. As these complex-phase modes propagate across the cortical surface, they interact: Where the phases constructively interfere, the result is synchronised activity, while where they destructively interfere, activity is dampened. These transient, localised zones of constructive interference are exactly what we call turbulent eddies in fluid dynamics. They are non-stationary packets of energy and information.

The deep reason why this unification matters lies in what it does for the three parts of the architecture of the Entangled Loop. In this theory, emotion is architecturally primary. Evolution’s response to the intractability of survival was not to compute better but to compress, and valence, the good-for-me or bad-for-me of an encountered state, is that compression (Kringelbach et al., 2026, 2024b). Emotion in this sense is not content that the conscious mind happens to represent; it is the boundary condition that shapes the attractor landscape of the world model. The empirical realisation of this compression used by evolution is described by the so-called ‘pleasure cycle’ of wanting, liking and satiety (Berridge and Kringelbach, 2008, 2015). This affective-primacy position for explaining consciousness has a long history through Panksepp, Solms, Damasio and Craig (Panksepp, 1998; Solms and Friston, 2018; Damasio, 1994; Craig, 2015), but the Entangled Loop is the first treatment to derive it from thermodynamic first principles.

In the brain the spectral gap of the connectome is produced by the rare long-range connections that violate the exponential distance rule and that are particularly enriched in humans (Markov et al., 2014; Deco et al., 2021a). The gap opens the interference regime, which Deco and colleagues have incorporated into a quantum-like whole-brain model that fits empirical neuroimaging significantly better than its classical counterpart at lower energy cost (Deco et al., 2025c). The inference carried on this substrate is then quantum-like in the precise sense that it performs everything classical Bayesian inference performs and additionally supports interference effects no finite classical circuit can reproduce (Busemeyer and Bruza, 2012; Pothos and Busemeyer, 2009; Aerts, 2009; Riechers et al., 2025; Kringelbach et al., 2026). The thermodynamic argument completes the picture, since a brain under selection pressure to compute more at lower energetic cost will converge on whichever regime supports the richest probability structure at the lowest dissipation, and that regime is the quantum-like one. Binding, in the sense of phenomenology, is the question of why experience is one perspectival field rather than scattered, and it falls out of this interference rather than being added on top of a feedforward stack. The theory’s empirical signature in this register is the Växjö Interference Connectivity, whose pre-clinical version predicts adolescent anxiety and depression at one-year follow-up substantially above the functional connectivity ceiling (Deco et al., 2026).

The second feature is that the architecture uses interference to support an inference regime that is quantum-like rather than classical Bayesian. The distinction matters and is easy to blur, so we state it carefully. As shown above, by *interference* we mean the physical phenomenon, in which coupled oscillators on a network with a spectral gap superpose with constructive and destructive amplitudes that do not reduce to additive probability (Scholes, 2024; Scholes and Amati, 2025). By *inference* we mean the computational use of the resulting state space to update beliefs about the world. The bridge between them is a theorem of Khrennikov and colleagues, the Växjö interpretation, which establishes that a classical system whose amplitudes interfere produces probability structure with the algebraic form of quantum mechanics, including contextuality and order effects, without any microscopic quantum substrate (Khrennikov, 2010; Bulinski and Khrennikov, 2004).

The third feature is hierarchical orchestration. Integration across spatial and temporal scales is not free: It demands coordination, and that coordination is the work of a small set of orchestrating regions sitting at the top of the cortical information flow (Deco et al., 2021b; Margulies et al., 2016; Glomb et al., 2021). The same regions emerge from the analysis of consciousness across cognitive tasks and from the analysis of brain states, so the orchestrating ensemble is one structure, not an ad hoc list (Mashour et al., 2020; Kringelbach et al., 2024b).

The three features are inseparable because they share a single thermodynamic ancestor. Emotion is the compression that makes survival affordable on a metabolic budget of about twenty watts, the interference is the binding that makes that compression a unified perspective, and the hierarchical orchestration is the coordination that holds the two together. The theory builds on the *Beautiful Loop* of Laukkonen et al. (2025), which treats consciousness as self-evidencing inference in a world model, and adds exactly these two elements, the architectural primacy of emotion and the quantum-like character of the binding inference, supported by the interference its substrate makes available. The word “entangled” is meant in the precise sense that emotion, interference and hierarchy cannot be separated, and “loop” in the sense that the architecture is self-evidencing.

The empirical test in Section 6 provides strong support for the single-operator scaffolding on which this architecture rests. Specifically, it sharpens the claim from an exact operator identity to a falsifiable prediction on the brain. Perturbing the operator by a 5-HT_2A_ receptor density map and asking one scalar coupling to predict, simultaneously, the multi-scale turbulence shift under LSD and the macroscale harmonic energy redistribution recovers a single value *α*^∗^ = 0.0060 at which both correlations are high (*R*_turb_ = 0.987, *R*_energy_ = 0.805). The two predictions live in mathematically independent domains, the spatial resolvent and the discrete spectral shift, and have no reason to share a maximum unless the underlying operator is one and the same. The spectral gap is preserved to 5 ×10^−4^, so the structural hierarchy is robust under the perturbation, while the multi-scale dynamics reorganise around it. These observations are the operator-calculus identity tested on the brain rather than asserted, and seam (S1) is empirically tolerable at the macroscale where the theory makes its claims.

We propose that this is the necessary mathematical scaffolding of the Entangled Loop. The hierarchy, the cascade and the interference are not three independent computational features of brain dynamics; they are three readings, in the strict sense of the spectral theorem, of one anatomical operator. The theory’s architectural claim about consciousness rests on this scaffolding, and the scaffolding is exact as operator calculus and approximate on the brain only to the extent that one explicit idealisation, the degree homogeneity of the connectome, fails. The LSD perturbation result is direct evidence that the degree-twist sits within tolerance at macroscale. Appendix B catalogues the seams in honest detail. The empirical reach of the unification beyond the LSD test is demonstrated in our companion preprint (Deco et al., 2026), where a signature of entanglement derived from the same operator is shown to carry predictive information about adolescent depression and anxiety at one-year follow-up.

The wider biological architecture of the Entangled Loop lies outside the scaffolding. The compressed valence that closes the loop is a separate mathematical object, and the affective-primacy claim that drives the theory is a claim about what selects *δV* in Equation (12), not about the spectral identity of Section 3. We close by stating the research programme this leaves open.

Reinforcement learning is the natural first place to look for a closure rule. We propose that it is not the right primitive for the Entangled Loop, for three structural reasons. First, reinforcement learning takes an exogenous scalar reward as given and learns to maximise its expected discounted return, whereas in the present theory valuation is itself the thermodynamic compression of the survival decision space and is therefore the object to be explained (Kringelbach et al., 2026, 2024a). Second, the reinforcement-learning objective is a fixed point to be approached, whereas survival is the maintenance of viability, a constraint and a Lyapunov structure rather than an unbounded maximum. Third, the reinforcement-learning return is acyclic and time-averaged, while the pleasure cycle is intrinsically a self-renewing transient in which wanting, liking and satiety follow one another and dissolve, the felt life of the organism rather than its asymptote (Berridge and Kringelbach, 2008, 2015; Kringelbach et al., 2024b). Reinforcement learning is recoverable as the flattened, fixed-point shadow of this cyclic structure, useful where the shadow is the part one needs to compute, but not the fundamental object.

We propose instead that the pleasure cycle is evolution’s solution to the same thermodynamic problem that selects the rest of the architecture. A homeostatic state must be held within a viability set at minimal energetic cost, and the cycle of wanting, liking and satiety is the affective dynamics that does this by being intrinsically unstable. Stability would mean the organism could rest at a single attractor, which is also the state of maladaptive narrowing that depression and addiction realise, the rigid low-dimensional landscape of the depressed brain (Perl et al., 2023; Pizzagalli, 2022; Husain and Roiser, 2018; Treadway and Zald, 2011; Rømer Thomsen et al., 2015). The healthy pleasure cycle is the limit cycle around viability, continually leaving and re-entering the set point, which keeps the operator continually re-tuned, keeps the dynamical repertoire broad and keeps the system close to the near-criticality at which best fit, maximal turbulence and maximal information capability coincide (Kringelbach et al., 2024a; Atasoy et al., 2019; Deco et al., 2025a). Instability and flexibility are the same property read twice. The cycle is unstable in the sense that it does not settle, and flexible in the sense that the system it carries can re-enter any state from the viable repertoire.

Flourishing, on this view, is not a steady state but the sustained operation of the cycle with the full hedonic repertoire intact and meaning-making integrated into it. It is meaningful pleasure in the sense of Kringelbach et al. (2024b), intrinsically unstable and continually renewed, which connects the present account to the functional neuroanatomy of pleasure and happiness developed in the Kringelbach laboratory (Kringelbach and Berridge, 2009; Berridge and Kringelbach, 2011). The mathematical form of this closure is not stated in the present paper. Plausibly it is a viability functional whose dynamics is a relaxation limit cycle, with species survival entering as a kin-weighted inclusive-viability term, and the slow control descending a Lyapunov gradient on the spectrum of the same operator the scaffolding identifies. The conjecture is the subject of a separate treatment. The point we close on here is that the scaffolding of Section 3 is what makes that closure intelligible, since the hand that re-tunes the operator is emotion, and emotion can be the architecturally primary boundary condition of the world model only because the operator is one operator. The mechanism of consciousness, in the sense of the architecture rather than the content, is one operator read three ways, and emotion is what selects the perturbation under which it is read.

## A The resolvent identification

This appendix gives the detailed derivation that the turbulence kernel is the resolvent of the harmonic operator. The argument has three steps, namely the identification of the exponential as a screened Green’s function, the spectral expansion of the resolvent, and the application of the functional calculus to recover the local order parameter in eigenbasis form. Lemma 1 and Proposition 2 contain the result in compressed form. We unpack them here.

### Step 1: The exponential as a Green’s function

The local order parameter of Equation (2) smooths the complex phase field *e*^*iφ*(*x,t*)^ by an exponential kernel *G*_*λ*_(*x*) = *e*^−*λ*|*x*|^. In the one-dimensional homogeneous continuum, this kernel is the Green’s function of the modified Helmholtz operator −*d*^2^*/dx*^2^ + *λ*^2^. To see this, take the Fourier transform of the equation (− Δ +*λ*^2^)*G* = *δ* to obtain 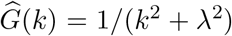, and invert with a contour integral closed in the lower half-plane,

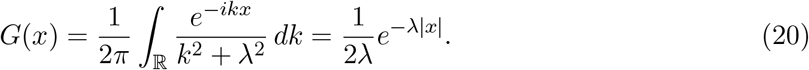

Equivalently, away from the origin *G* satisfies *G*^′′^ = *λ*^2^*G* with the decaying solutions *e*^∓*λx*^. Continuity at the origin and the unit jump *G*^′^(0^+^) −*G*^′^(0^−^) = −1 fix the amplitude 1*/*(2*λ*). In three dimensions the same argument gives the Yukawa potential *G*(*r*) = *e*^−*λr*^*/*(4*πr*).

The empirical kernel of the turbulence model and the empirical kernel of the anatomy, namely the exponential distance rule, are therefore not two different phenomenological choices that happen to have the same form. They are the same mathematical object, the screened Green’s function, evaluated either as a kernel on the cortical surface (the EDR) or as a smoothing applied to a field defined on it (the local order parameter).

#### Step 2: The resolvent in eigenbasis

Let *L* have spectral resolution *L* = ∑_*k*_ *µ*_*k*_*P*_*k*_ with 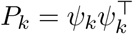. The resolvent at −*λ*^2^ is defined by the functional calculus

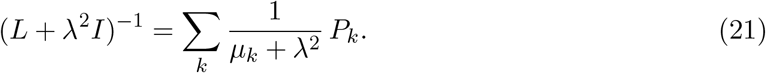

This is Proposition 1 with *f* (*µ*) = (*µ* + *λ*^2^)^−1^. The weights 1*/*(*µ*_*k*_ + *λ*^2^) are monotone decreasing in *µ*_*k*_, so the resolvent acts as a low-pass filter in the eigenbasis, attenuating the high modes by the bulk size of their eigenvalues and preserving the low modes relatively undamped.

#### Step 3: Recovering the local order parameter

The convolution form of Equation (2),

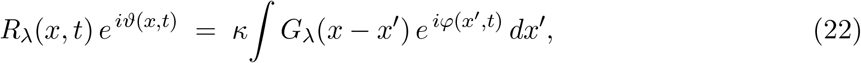

is the application of the convolution operator *G*_*λ*_∗ to the complex phase field. By Step 1, this convolution is, up to the constant 2*λ*, the inverse of the screened Laplacian, that is, *G*_*λ*_∗ = (2*λ*)^−1^(− Δ +*λ*^2^)^−1^. Expanding the phase field in the eigenbasis of − Δ,

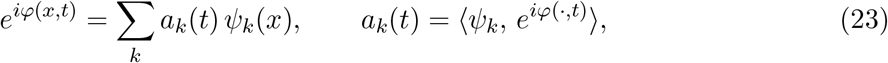

and applying the resolvent term by term,

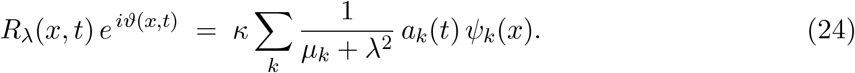

The local order parameter at time *t* is therefore the spectral expansion of *e*^*iφ*^ with the high modes attenuated by the resolvent weights. Its amplitude |*R*_*λ*_| and its spacetime variability *D*_*λ*_ are statistics of this expansion. The harmonic spine is the support, and the turbulent cascade is the dynamics of the modal amplitudes *a*_*k*_(*t*) filtered through the same support. The two are inseparable because they are the projection and the resolvent of one operator.

#### What the rare long-range exceptions do

Adding a few long edges to an EDR graph perturbs *W* by a low-rank update of large entries. By the Courant–Fischer eigenvalue inequalities this raises the algebraic connectivity *µ*_2_, hence the gap *g* = *µ*_2_ − *µ*_1_, more than it raises the bulk eigenvalues. The resolvent weight on the lowest non-trivial mode 1*/*(*µ*_2_ + *λ*^2^) therefore decreases relative to the bulk, which sharpens the low-rank dominance of the resolvent. Geometrically, this is why the EDR+LR connectome, which the literature has identified as enhancing the information cascade, also gives the cleanest spectral hierarchy. The mechanism is one and the same.

#### Hierarchy as the low-rank skeleton, restated

The first nontrivial functional harmonics coincide with the principal cortical gradient and therefore with the cortical hierarchy (Glomb et al., 2021; Margulies et al., 2016). By Proposition 2 the resolvent is dominated by these same low modes once the gap is large enough. The hierarchy is therefore not merely correlated with the turbulence kernel; it is the low-rank skeleton of the resolvent and so the low-rank skeleton of the turbulence kernel. The empirical fact that the orchestrating regions sit at the unimodal-to-transmodal extreme of the principal gradient (Margulies et al., 2016; Deco et al., 2021b) is the same statement read on the brain: The orchestrating regions are the regions that load on the resolvent’s dominant modes, which is what gives them their disproportionate effect on global dynamics.

#### CHARM as the propagator on the same skeleton

The Schrödinger factor 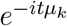 rotates each *ψ*_*k*_ by a phase 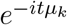 per unit time. A gap separates the angular velocities *µ*_*k*_ of the low modes from the dense angular velocities of the bulk. The interference pattern carried by the sub-gap modes is therefore low-dimensional and structured, and it lives on exactly the eigenbasis the harmonics name and the resolvent dominates. The CHARM form measures this structured interference, and the canary’s Växjö Interference Connectivity reads it through the violation of the law of total probability in vortex space (Deco et al., 2026, 2025b). The functional calculus places the harmonic skeleton, the turbulence kernel and the CHARM form on one operator and one set of eigenvalues, and the gap moves all three together.

#### B The seams: Exactly where this can fail

The unification is exact at the level of functional calculus (Proposition 1). Its application to the brain rests on identifications that are clean only in idealisations, and these are the points to attack. We catalogue them honestly.

### (S1) Graph Laplacian versus screened-Laplacian resolvent. The load-bearing seam

Proposition 2 shows the EDR kernel is the resolvent of a continuum screened Laplacian Δ_0_, so that *W* ≈ (− Δ_0_ +*λ*^2^)^−1^ sampled on the nodes. But connectome harmonics are eigenvectors of *L* = *D* − *W*, not of *W* or of Δ_0_. If *W* = (− Δ_0_ +*λ*^2^)^−1^ then *W* and Δ_0_ share eigenvectors, yet *L* = *D* − *W* shares them only if *D* commutes with *W*, that is, if the degree is constant 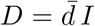 (a degree-homogeneous graph). With heterogeneous degree, [*D, W*] ≠ 0 and the connectome harmonics are the eigenvectors of *D* − *W*, a degree-twisted version of the Δ_0_ harmonics. The identity that harmonics are eigenfunctions of the operator whose resolvent is the EDR kernel is therefore exact only under degree homogeneity, and is otherwise a controlled perturbation governed by the degree variation. The normalised graph Laplacian *I* − *D*^−1*/*2^*WD*^−1*/*2^ mitigates but does not remove this. Quantitatively the mixing is governed by ∥[*D, W*]∥ relative to the gap *g*, and the synthesis is safe when the degree spread is small compared with *g*. The whole load of the present identification on the brain rests here.

### (S2) Euclidean versus intrinsic distance

The EDR uses Euclidean *r*(*n, p*). The harmonic operator’s natural metric is the graph or manifold geodesic. Lemma 1 is a flat-space statement, and on the folded cortex Euclidean and geodesic distances differ, so the identification of *W* with the resolvent is metric-approximate.

### (S3) Two distinct appearances of *e*^−*λ*|·|^

The exponential appears twice, as the dynamical coupling *C*_*np*_ in the Stuart–Landau equations and as the observational smoothing *G*_*λ*_ in the definition of *R*_*λ*_. Proposition 2 concerns the observational kernel. That the dynamical coupling is governed by the same operator requires the linearisation of (S4), and the two *λ* need not be equal.

### (S4) Linear spectral picture versus nonlinear turbulence

The functional calculus is linear. Stuart–Landau and Hopf turbulence are nonlinear. The unification is exact for the linear backbone, that is, the Hopf normal form linearised at the operating point with its linear part the connectome-coupled operator, and for the observational resolvent. It is approximate for fully developed nonlinear turbulence. The empirical coincidence of best fit, maximal turbulence and near-criticality at a single coupling (Deco et al., 2025a) is consistent with the linear backbone being the organising skeleton, while the cascade across the inertial subrange is a nonlinear phenomenon the spectral identity alone does not deliver.

### (S5) Dimension

The function *e*^−*λ*|*x*|^ is the exact one-dimensional screened Green’s function. In three dimensions it is *e*^−*λr*^*/*(4*πr*). The EDR is stated empirically as a pure exponential. The cleanest exactness is therefore on a one-dimensional ring model, which the turbulence literature uses explicitly, and is an approximation in three-dimensional cortex.

### (S6) Wick rotation, formal versus physical

The map *e*^−*tL*^ → *e*^−*itL*^ is exact as functional calculus, since 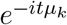 is defined on the spectrum, changing a contractive semigroup into a unitary group. The physical reading that the brain “uses” the Schrödinger kernel is a representational choice explicitly disclaimed as non-quantum in the CHARM paper (Deco et al., 2025b). The operator-calculus identity stands regardless of that reading.

### What is load-bearing

The load-bearing claim is Proposition 1 together with Proposition 2, namely that harmonics, the turbulence smoothing kernel and the CHARM form are one operator’s spectral measure read through the projection, the resolvent and the unitary propagator. As pure operator calculus this is exact. Its purchase on the brain stands or falls on seam (S1), namely whether the degree heterogeneity in the connectome is small enough relative to the spectral gap that the eigenvectors of *D* − *W* remain close to those of the screened Laplacian whose resolvent is the EDR kernel. If (S1) is tolerable, the harmonic spine and the turbulent cascade are literally one spine and the unification is real. If (S1) is not tolerable, the relationship is an analogy of form, not an identity, and must be stated in those weaker terms. Everything else in the note is either textbook or established in the primary papers cited. The empirical question of whether the human connectome is sufficiently degree-homogeneous, on the scale set by the spectral gap, is the single question that decides the strength of the synthesis.

## Acknowledgements

G.D. is supported by Grant PID2022-136216NB-I00 funded by MICIU/AEI/10.13039/501100011033 and by “ERDF A way of making Europe”, ERDF, EU, Project NEurological MEchanismS of Injury, and Sleep-like cellular dynamics (NEMESIS) (ref. 101071900) funded by the EU ERC Synergy Horizon Europe, and AGAUR research support grant (ref. 2021 SGR 00917) funded by the Department of Research and Universities of the Generalitat of Catalunya. M.L.K. is supported by the Centre for Eudaimonia and Human Flourishing (funded by the Pettit and Carlsberg Foundations) and Center for Music in the Brain (funded by the Danish National Research Foundation, DNRF117). The funders had no role in study design, data collection and analysis, decision to publish or preparation of the manuscript. The authors used Claude (Anthropic) for mathematical formalisation, structural editing and reference verification. All scientific claims are the authors’ own.

## Author contributions

Following the CRediT taxonomy: M.L.K. and G.D. both contributed to conceptualisation, methodology, formal analysis, investigation, software, data curation, validation, project administration, supervision, writing – original draft, and writing – review and editing.

## Code availability

All code used to reproduce the analyses reported in this paper, including the perturbative forward-model of Section 6, is openly available at https://github.com/decolab-public/unification.

